# A Leak-Free Head-Out Plethysmography System to Accurately Assess Lung Function in Mice

**DOI:** 10.1101/2021.12.01.470843

**Authors:** Stephanie Bruggink, Kyle Kentch, Jason Kronenfeld, Benjamin J. Renquist

## Abstract

Mice are a valuable model for elegant studies of complex, systems-dependent diseases, including pulmonary diseases. Current tools to assess lung function in mice are either terminal or lack accuracy. We set out to develop a low-cost, accurate, head-out variable-pressure plethysmography system to allow for repeated, non-terminal measurements of lung function in mice. Current head-out plethysmography systems are limited by air leaks that prevent accurate measures of volume and flow. We designed an inflatable cuff that encompasses the mouse’s neck preventing air leak. We wrote corresponding software to collect and analyze the data, remove movement artifacts, and automatically calibrate each dataset. This software calculates inspiratory/expiratory volume, inspiratory/expiratory time, breaths per minute, enhanced pause, mid-expiratory flow, and end-inspiratory pause. To validate the use, we established our plethysmography system accurately measured tidal breathing, the bronchoconstrictive response to methacholine, sex and age associated changes in breathing, and breathing changes associated with house dust mite sensitization. Our estimates of volume, flow, and timing of breaths are in line with published estimates, we observed dose-dependent decreases in volume and flow in response to methacholine (P < 0.05), increased lung volume and decreased breathing rate with aging (P < 0.05), and that house dust mite sensitization decreased tidal volume and flow (P <0.05) while exacerbating the methacholine induced increases in inspiratory and expiratory time (P < 0.05). We describe an accurate, sensitive, low-cost, head-out plethysmography system that allows for longitudinal studies of pulmonary disease in mice.

**New & Noteworthy:** We describe a variable-pressure head-out plethysmography system that can be used to assess lung function in mice. A balloon cuff that inflates around the mouse’s neck prevents air leak, allowing for accurate measurements of lung volume and air flow. Custom software facilitates system calibration, removes movement artifacts, and eases data analysis. The system was validated by measuring tidal breathing, responses to methacholine, and changes associated with house dust mite sensitization, sex, and aging.

**Contributions to Study:**

1. Stephanie Bruggink: development of head-out plethysmography chamber, measurement of breathing, data analysis, prepared manuscript
2. Kyle Kentch: development of head-out plethysmography chamber, programmed software to collect and analyze data, prepared manuscript
3. Jason Kronenfeld: development of tools to analyze data, analysis of data
4. Benjamin Renquist: development of head-out plethysmography chamber, statistical analysis, prepared manuscript

## Introduction

Mice are a valuable experimental model to study complex, systems-dependent diseases. However, studies of the pulmonary system in mice are limited by the tools available to assess lung function. We set out to develop a low-cost, accurate, head-out variable-pressure plethysmography system to allow for repeated, sensitive, non-terminal measurements of lung function in mice. Common methods to assess lung function in mice include whole-body plethysmography, head-out plethysmography, and forced oscillation technique. However, these popular techniques have limitations.

With whole-body plethysmography the mouse is placed in a chamber large enough to allow for unrestricted movement, and changes in chamber pressure are measured. Changes in pressure or flow are used to assess lung volume and result from both alterations in air temperature and humidity as air enters/leaves the lung and alveolar gas compression and decompression (1-3). More than one-half of the change in chamber pressure or flow associated with breathing can be eliminated by equilibrating the humidity and temperature within the chamber with mouse’s body temperature and the humidity of exhaled air (4). Thus, unless the chamber is set to mirror the mouse’s body temperature and the humidity of exhaled air, the measured change in pressure or flow is partially an experimental artifact that cannot be quantified. This is a difficult challenge in a system that relies on continuous flow of air through the chamber (5). The remaining signal is a measure of alveolar compression and decompression of air, associated with flow and volume, two measures affected by airway resistance (4, 6).

Forced oscillation technique measures the impedance of gas waveforms directed into the lung of an anesthetized, paralyzed, tracheostomized, and ventilated mouse (7). This technique is sensitive, creating reproducible measures of airway mechanics. Different oscillation maneuvers can be applied to determine resistance, compliance, elastance, input impedance, and the energy dissipated or conserved within the lung parenchyma (tissue damping/elastance). Resistance describes the patency of the conducting airway (resistance of the airway to airflow) and compliance describes the ability of lungs to expand (8, 9). This technique is precise, but, accuracy requires calibration, limiting resistance of the endotracheal cannula, proper positioning of the animal, standardization of lung volume history, and a passive respiratory system (10). Standardization is necessary because oscillations of different frequencies and volumes alter respiratory mechanics including resistance and compliance. Controlling the waveform oscillation with the same software program and technique for each measurement can resolve variability (2, 11-13). However, even when these technical challenges are overcome, this technique is limited. It is terminal, preventing repeated measures across time in the same mouse, and, it cannot be applied to understand alterations in the respiratory drive of breathing as mice are anesthetized and paralyzed (2).

Orotracheally intubating anesthetized mice is described as an alternative to forced oscillation technique because it allows for non-terminal measurements in spontaneously breathing animals (14). Flow is assessed from the orotracheal tube and transpulmonary pressure is assessed by measuring pressure changes from a water-filled esophageal tube (15). Still, this measurement must be performed in the anesthetized mouse. Since normal breathing patterns are altered with anesthesia, we aimed to identify a system that measured breathing in awake, conscious mice (16). Head-out plethysmography assesses lung function by measuring flow or pressure changes inside a chamber caused by expansion and contraction of the ribcage as the animal inhales and exhales. Unlike whole-body plethysmography, in which the pressure change that results from ribcage expansion is negated by the concomitant removal of chamber air with inhalation, with head-out plethysmography the inhaled air is coming from outside the chamber. In turn, the pressure change resulting from expansion of the ribcage is not negated by a simultaneous removal of air from the chamber (17). Unlike the forced oscillation technique, head-out plethysmography is non-terminal and performed in awake, conscious mice. In turn, head-out plethysmography allows for repeated measurements of physiologically relevant breathing. All published and commercial plethysmography chambers use a piece of latex (or other flexible plastic) to create an airtight seal around the mouse’s neck (17). In preliminary studies we found this set-up did not create an airtight seal. Without an airtight seal, pressure-volume relationships to determine tidal volume and flow are inaccurate. Accordingly, as described, head-out plethysmography can provide accurate measures of the timing of breaths but cannot be applied to accurately assess either volume or flow.

Double-chamber plethysmography utilizes a similar set-up to head-out plethysmography with an additional chamber fit around the head or nose of the mouse. Pressure changes resulting from nasal flow and thoracic movements are measured separately (18). Importantly, this allows for measurement of specific airway resistance, assessed by the time delay between the nasal and thoracic flows. The change in lung volume proceeds the change in nasal flow, creating a pressure gradient to pull air into the lung. The delay between thoracic and nasal flows is short in healthy animals and increases with increased resistance. Although this dual chamber does have the added benefit of assessing resistance, accuracy is again limited by air leak between the head and body chambers (2, 14).

The lack of an accurate and sensitive tool to repeatedly measure breathing in the same mouse across time limits study design and mouse models that can be used to understand pulmonary disorders. We set out to create a leak-proof variable-pressure head-out plethysmography system that would allow for accurate and sensitive measurements of volume and flow. Because this system records data corresponding to each breath, we created software to easily collect and manage the large amounts of data obtained from each mouse.

## Materials & Methods

### Head-Out Plethysmography Chamber

The main body of the variable-pressure plethysmography double chamber is composed of two syringe barrels (inner chamber 60 ml Monoject #1186000555; outer chamber 300 ml A AKRAF #B07T7MN36N) with the hollow tip end removed from larger barrel (Figure 1.7). The larger syringe is cut shorter allowing the smaller syringe to extend beyond the larger syringe on both ends (Figure 1.7). The smaller syringe is held in place within the larger syringe using 3D-printed spacers at both ends (Figure 1I, 1.2). O-rings that fit tightly into grooves on the printed spacers create an airtight seal where the spacers meet the syringe walls (Figure 1F inner O-rings Danco #35736B 1.25 inch diameter x 1 inch diameter; Figure 1G outer O-rings Danco #35763B 1.88 inch diameter x 1.62 inch diameter). The spacers must be fit onto the smaller syringe before removing the hollow tip of the inner syringe (Figure 1.2). A small amount of petroleum jelly may facilitate this. Next, the smaller syringe is perforated and hollow tip end removed (Figure 1.5). The perforations allow air to freely flow from the inner chamber to the outer chamber. To prevent air-leak, all perforations on the inner chamber must be between the two spacers. This double chamber system prevents the mouse from occluding the air input and pressure gauge attached to the outer syringe. The syringe plunger (Figure 1A) creates an air-tight seal at the back of the chamber. At the front of the inner chamber is an inflatable balloon cuff that allows the user to create an airtight seal around the mouse’s neck. This balloon cuff is made from a 3D-printed ring (Figure 1J; 3-D Printer File at https://github.com/bjrenquist/plethysmography/tree/main/printed_parts) with the midline lined with holes that allow air to pass from the outside to inside of the ring to inflate the balloon cuff. Both edges of the ring have groves in which O-rings sit (Figure 1E Danco O-rings #35731B 0.94 inch diameter x 0.75 inch diameter). To affix the balloon, a small latex balloon (Figure 1K Amscan #115914) is cut into a ∼2 cm wide ring and put inside of the 3-D printed ring (Figure 1.3). The balloon ends are wrapped over the outside of the ring and held in place with O-rings (Figure 1.6). To allow inflation of the balloon, we use a needle (any size and brand) to perforate the balloon where the balloon ends overlap and occlude the holes along the midline of this ring (Figure 1.6).

**Figure 1.**
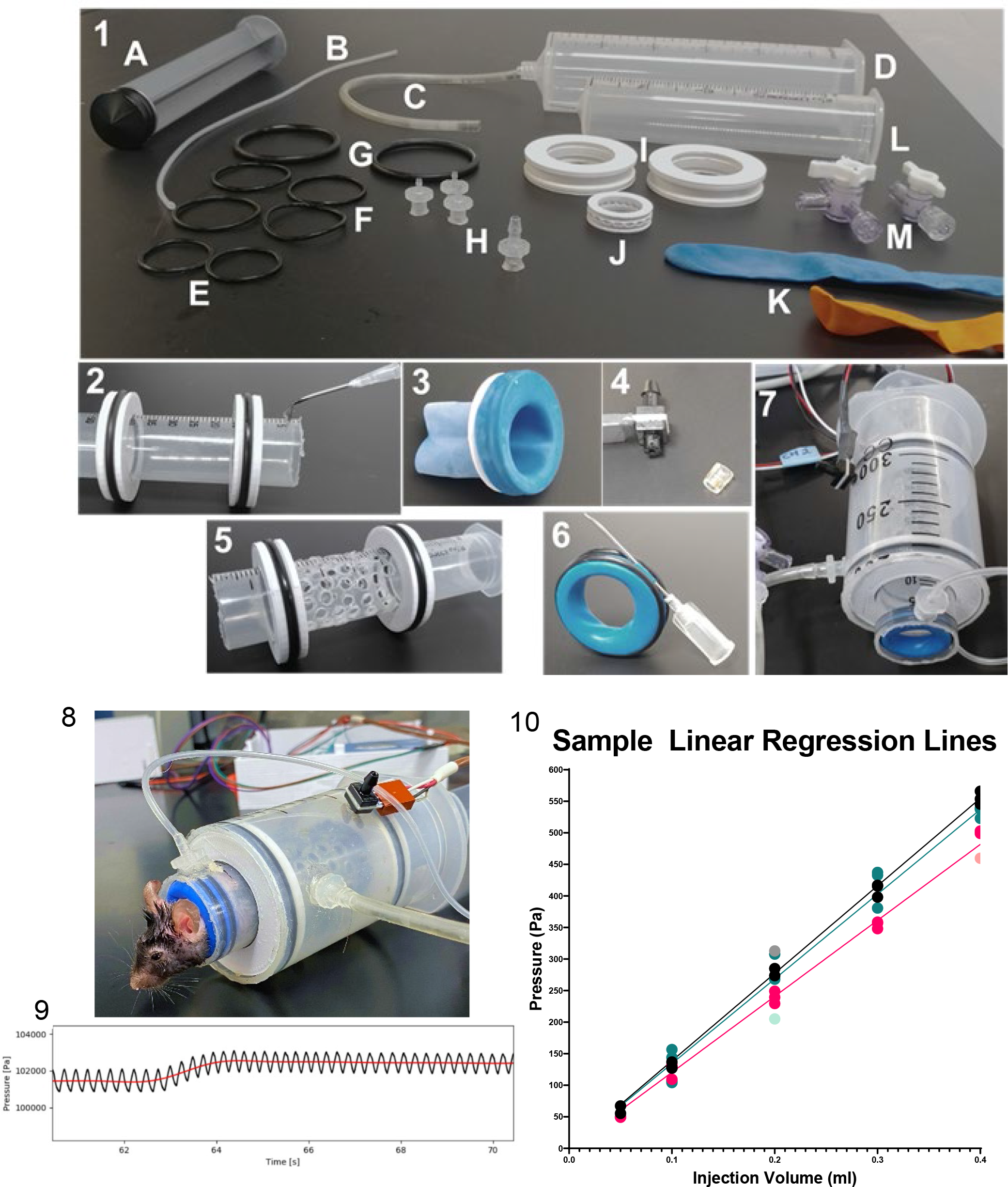
Construction of a leak proof, variable-pressure head-out plethysmography chamber that can be calibrated. 1) Head-out plethysmography chamber materials. A. Plunger fitting in 60 ml syringe (L) to prevent mouse from backing out of chamber B. Tubing to create balloon cuff air port C. Tubing to create calibration air port D. 300 ml outer syringe E. Two O-rings to secure balloon to 3-D printed ring for balloon cuff (J) F. Four inner O-rings to secure 3D printed spacer (into I) G. Two outer O-rings to secure 3D printed spacer (into I) H. Female luers to create air port and attach tubing to stopcocks (M) I. Two 3D printed spacers J. 3D printed ring for balloon cuff K. Two options for balloons to create balloon cuff L. Inner chamber 60 ml syringe M. Stopcocks to allow air flow into balloon cuff and out chamber 2) Inner chamber with 3D printed spacers and O-rings fitted into grooves, picturing creation of hole for balloon port 3) Preparation of balloon cuff to be placed directly under hole (created in 2) 4) Pressure transducer with tip removed from one end and small piece of plastic tubing to place over the tip 5) Inner chamber complete with holes for adequate assessment of pressure changes by pressure transducer 6) Sample of how to perforate balloon cuff to allow inflation 7) Complete set-up with inner and outer chamber, balloon cuff, pressure ransducer, and ports to allow for airflow into the chamber or into the balloon 8) mouse in plethysmography chamber 9) Sample pressure trace with breaths signified in black and the red line indicating average pressure which shifts after injection of a calibrating quantity of air at 63 seconds. 10) Sample of calibration regression lines created for 3 mice (pink, marine, and black) with automatically excluded points in corresponding lighter color.

We created a small air port (hole created with needle) in the inner syringe barrel to inflate the balloon cuff (Figure 1.2). Next, we placed the outer chamber over the inner chamber making sure not to move the 3D printed spacers, a process facilitated with a small amount of petroleum jelly. A female luer (Figure 1H thread style to 500 series barb 1/16 inch ID tubing Cole-Parmer Instrument #SK-45508-01 connected to tubing Picture 1B Silastic Laboratory Tubing #508-005) was briefly heated to melt the plastic and this melted leur was placed over the top of the hole and allowed to cool and adhere creating an air port onto which tubing could be attached. The tubing, cut to ∼22 cm, is then connected to another female luer and a stopcock (Figure 1M Cole-Parmer #30600-00). This allows for the balloon cuff on the inside of the inner chamber to be inflated by a 3 ml syringe from the stopcock.

A small hole (0.5 centimeter in diameter) in the outer chamber fits the barbed end of a dual barbed pressure transducer (Honeywell SSCSAAN004NDAA5 board mount pressure sensor SIP, dual ax barbed differential, 5 Volts), surrounded by a small segment of flexible tubing (Tygon #R-3603) to allow for an air-tight seal (Figure 1.4). The second barbed port of the pressure transducer is left open to atmospheric air. On the outer chamber another air port, created by placing a female luer over a small hole, is used for calibration (Figure 1H Biorad #7318223 connected to tubing Picture 1C Tygon #R-3603). A stopcock (Figure 1M Masterflex #30600-02) is connected to the calibration tubing (cut to ∼22 centimeters) via another female luer and air is injected with any 1 ml syringe (Figure 1.7). A data acquisition unit (Measurement Computing, USB-1208FS-Plus) controlled by LabVIEW collects the voltage signal output from the pressure transducer. Multiple chambers and pressure transducers can be connected to the data acquisition unit to allow for the user to simultaneously measure breathing in multiple mice.

### Mouse Preparation and Loading into Chamber

Prior to starting breathing measurements the mouse’s neck is shaved and the remaining hair removed using a depilatory. This provides a smooth skin surface interfacing with the balloon cuff allowing for an airtight seal.

Mice are scruffed and gently placed headfirst in the plethysmography chamber. Once entirely in the chamber, the back is occluded with a syringe plunger and the mouse is coaxed toward the front of the chamber with the plunger. To prevent the mouse from biting and puncturing the balloon during loading, a small, flexible piece of plastic is set on the balloon and removed after the mouse has extended its head past the cuff and the plunger is far enough forward so the mouse cannot pull its head back into the chamber. Once the mouse has extended their head beyond the balloon cuff, a syringe, attached to the balloon air port, inflates the balloon restricting forward movement (Figure 1.8). To ensure an airtight seal, we use a 6 mL syringe filled with petroleum jelly to place petroleum jelly at the interface of the neck and balloon. Prior to each measurement we confirm the chamber is airtight by removing or adding air through the calibration port and verifying the induced change in pressure holds steady at this new baseline (Figure 1.9).

### Volume Calibration

Since each mouse displaces different chamber volumes, the user must calibrate the volume-pressure relationship prior to each breathing measurement. We calibrate by rapidly injecting known volumes of air that range from 0.05-0.4 ml (0.05, 0.1, 0.2, 0.3, and 0.4 ml), as the inspiratory and expiratory volume is expected to be < 0.4 ml. We then quantify the pressure change associated with injection of a given volume. The chamber pressure is reset to atmospheric pressure by opening the 3-way stopcock, or simply removing the syringe from the stopcock, allowing room air to flow into the air port between each injection. To ensure precise calibration, each volume is injected 4 times. The software has a built-in annotation marker, allowing us to log the injection volume and timing (Figure 2A, C). Automated calibration software (use described below) allows the user to enumerate the pressure-volume relationship by regressing the change in pressure on volume injected and fitting an equation that will be used to extrapolate all changes in chamber pressure to changes in chamber volume associated with breathing (Figure 1.10).

**Figure 2.**
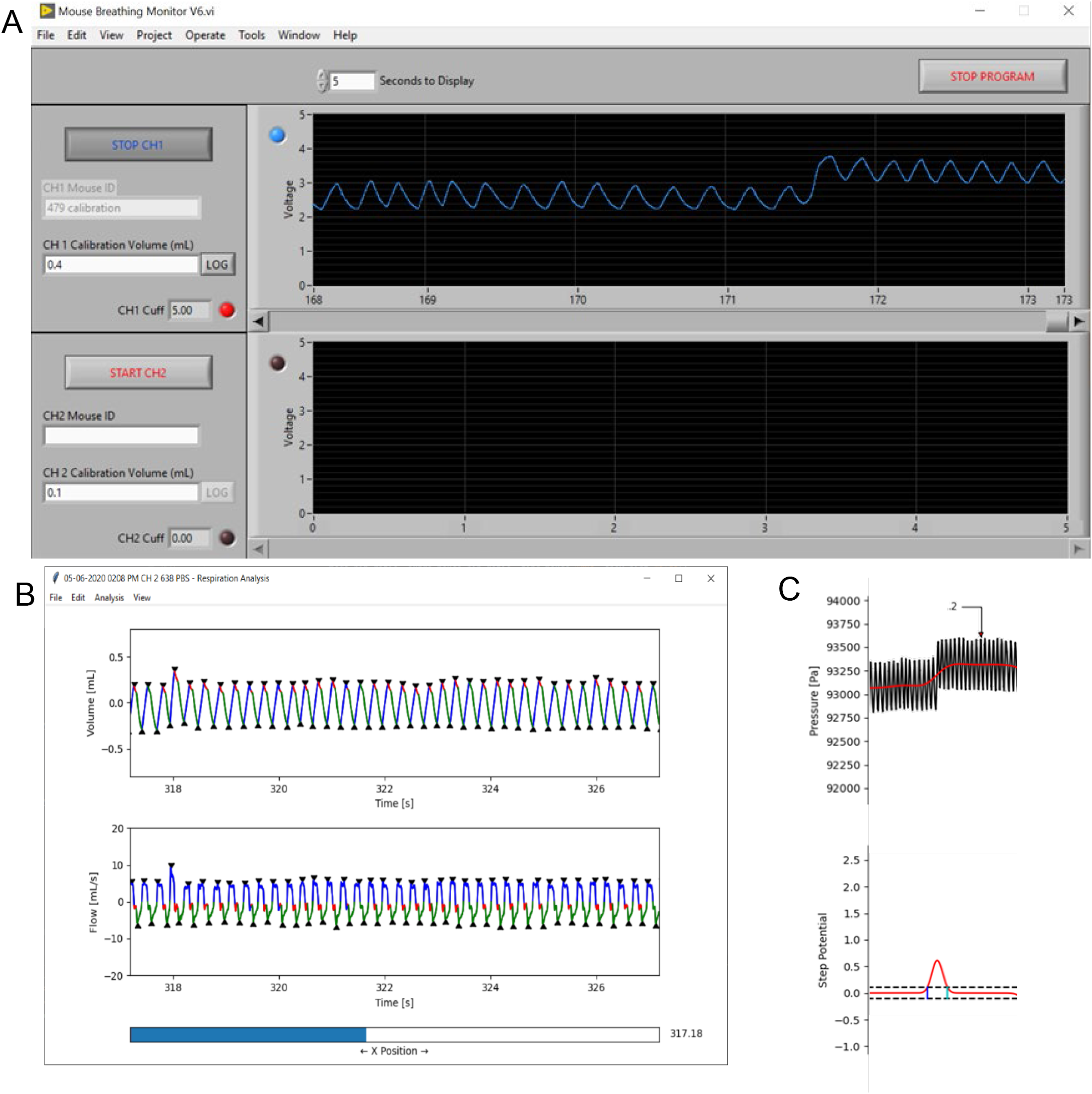
User interface for data collection and analysis software. A) Representative user interface for breathing data collection program. White boxes represent area where user can fill in data for mice from two separate chambers. B) Sample breathing trace opened with Python Respiratory Analysis Software with pressure converted to volume and flow and identified breath landmarks. Blue=inspiration, red=end-inspiratory pause, green=exhalation. C) Mouse breathing pressure signal during a 0.2ml calibration injection. Calibration injection signal, red, obtained after smoothing the pressure signal with a Gaussian filter (sigma=400 datapoints; top). Step potential, *θ_i_*, red, for the region of interest surrounding the calibration injection (bottom). Positive and negative step height thresholds, dotted black, set at 0.25 standard deviations from *θ_i_* = 0. Data points in peaks exceeding the step height thresholds represent step regions in the breathing signal. Data points within the step height thresholds represent plateau regions.

### Breathing Measurements

After calibration, breathing measurements are initiated. We waited at least 5 minutes for the mice to become accustomed to the chamber before collecting data, which was about the time it took to perform the calibration. When collecting only baseline measurements, we collected 10 minutes of tidal breathing. In studies to understand the response to methacholine, we collected a 5-minute baseline measurement followed by exposure to nebulized methacholine (MP Biomedicals, LLC). As multiple measurements were completed at different concentrations of methacholine, we averaged the 5-minute baseline measurements for each mouse to report tidal breathing. We exposed the mice to methacholine by placing the end of the neubulizer (Aeroneb lab nebulizer small unit, Kent Scientific) directly over the mouse’s head to ensure delivery into the airway. We delivered methacholine (dissolved in PBS) for 30 seconds at 0.191 ml/30 seconds at concentrations of 0, 25, 50, and 100 mg/ml of methacholine. Breathing was then measured for 10 minutes after the conclusion of methacholine administration. When completed, the balloon cuff was deflated then the syringe plunger was removed allowing the mouse to back out of the chamber. Mice were allowed to fully recover after each dose of methacholine by allowing at least 1 hour between measurements. Time course data is presented as 1-minute rolling averages taken every 30 seconds, while baseline data is presented as a 5-minute average. When applicable, we normalized to baseline values and present the data as change from baseline.

### House Dust Mite Sensitization

After completion of the dose response study, lean mice were sensitized using house dust mite (HDM). HDM (100 μg in 50 μl sterile PBS; D. Pteronyssinus; Stallergenes Greer; #XPB82D3A2.5) was administered intranasally on days 0, 7, and 14 (19). 24-hours after the last HDM administration, assessment of tidal breathing and response to methacholine was completed as described above.

### Data Collection Using LabVIEW

We developed a LabVIEW virtual instrument to facilitate the acquisition of real-time breathing measurements from up to two mice simultaneously. A set of driver software (Measurement Computing, ULx for NI labVIEW^TM^, Norton, MA; https://www.mccdaq.com/daq-software/universal-library-extensions-lv.aspx) was used to interface between LabVIEW and the data acquisition unit. This program samples the output voltage from each pressure transducer at a rate of 1000 Hz. The virtual instrument graphically displays the voltage signals in real-time for the user to visualize changes in pressure (Figure 2A). Titles and event timestamps allow the user to record an identifier for the mouse being studied and the volume and timing of calibration injections.

The user interface for data collection is simple (Figure 2A). Upon opening the program, the identifier for the mouse and measurement identifier was entered into CH1 or CH2 Mouse ID and measurement initiated by selecting *Start CH1/CH2*. For calibration, the volume injected was written into CH1/CH2 Calibration Volume, the volume was injected, and *LOG* was selected. Selecting *LOG* marked that timepoint with the entered volume (shown in Figure 2C). Of note, this event marker cannot be visualized in real time and only numbers can be entered. The *Start/Stop CH1/2* was used to begin and end each measurement on each separate channel. CH1/2 Cuff indicates the pressure of a second pressure transducer attached to the balloon cuff air port line allowing us to ensure we inflate the balloon to similar pressures for each measurement.

### Data Analysis Using Python Graphical User Interface

We developed a graphical user interface using Python 3.7 to convert the voltage signal, provided by the pressure transducer, into changes in chamber volume. Changes in chamber volume are representative of the mouse’s tidal volume and allow for calculation of expiratory and inspiratory volume, expiratory and inspiratory time, breaths per minute, enhanced pause (Penh), mid-expiratory flow (EF50), and end-inspiratory pause (Figure 2B).

The voltage signal from the pressure transducer is converted to a pressure signal based on the transducer’s specifications (Honeywell SSCSAAN004NDAA5) and its transfer function by *Equation 1*:

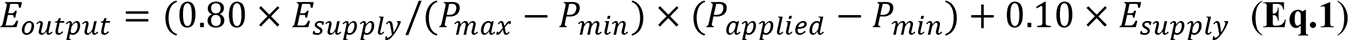

where *E_output_* is the output voltage signal, *E_supply_* is the voltage supplied to the pressure transducer, *P_max_* and *P_min_* are the maximum and minimum of the transducer’s pressure range, respectively, and *P_applied_* is the differential pressure between the measurement and reference ports of the transducer.

Calibration injections are used to empirically transform the pressure signal into a chamber volume signal. Air injected into the chamber creates a stable deflection in the pressure signaling which can be used to create a pressure-volume relationship (Figure 1.9, 1.10). To filter out the breathing signal and isolate the calibration injection signal, the pressure signal is smoothed using a Gaussian filter with a sigma of 400 datapoints (Figure 2C). For each calibration injection, a region of interest is initially bounded by the 24 seconds before and 12 seconds after the injection timepoint. Each region of interest is then refined by examining the local signal variance within two breathing periods (400 ms) of each datapoint. The final region of interest consists of a continuous segment of datapoints surrounding the injection timepoint where the local signal variance is no less than half the median value. This final region of interest is down sampled to improve processing speed, retaining 1 in every 4 datapoints.

To the smoothed pressure signal, we then applied a moving step fit algorithm in each region of interest (20). The published method was modified to optimize volume-pressure correlation and use the most datapoints to develop this correlation. To account for differences in signal-to-noise ratio, the half-window size, *w*, varies with calibration injection volume, *v*, such that *w* = 200 ⁄ *v*. Instead of a first-degree polynomial, we use a zero-degree polynomial for the fitting functions, under the assumption that the slope before and after a calibration step is zero. After obtaining *θ_i_* (the indicator for the probability of datapoint, *i*, to be a potential step position) a step height threshold is selected at 0.25 standard deviations from *θ_i_* = 0. Datapoints outside this threshold represent step regions and datapoints within the threshold represent plateau regions (Figure 2C). To be used in the analysis, a region of interest must consist of an initial baseline plateau followed by an upward step to a calibration plateau containing the calibration injection timepoint. Furthermore, each plateau must exceed a step width threshold of 0.5 seconds. The change in pressure for a calibration injection is calculated as the difference between the mean unsmoothed pressure signal values of the calibration and baseline plateaus.

The pressure changes of the calibration injections were matched with known injection volumes and a linear regression was used to determine the relationship between pressure and volume (Figure 1.10). Where *V_injection_* is the injection volume, and *m* is the slope. Given that there is no change in pressure without a change in volume, we set the *ΔP*-intercept (y-intercept) to 0.

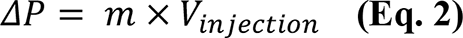

To ensure accuracy of the calibration, all injection volumes were repeated in quadruplicate and any points with a residual value exceeding one standard deviation away from the mean residual value were excluded.

A calibration injection causes an equivalent decrease in chamber volume (*V_injection_= -ΔV_chamber_*). Injection volume is substituted by change in chamber volume to calculate the mouse’s respiration signal:

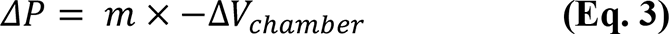

For our analyses, we report change in plethysmography chamber volume as a proxy for changes lung volume.

We used a Gaussian filter (sigma of 5 datapoints) to reduce noise present in the volume signal. This signal was then detrended to correct for any global drift in chamber pressure by fitting a line to the entire signal and subtracting this line from the signal. Further local drift correction was performed by subtracting the result of a two-second sliding average window from the global corrected volume signal. This correction for global and local drift helped to eliminate broad deflections in the breathing signals that were caused by small changes in chamber volume from the mouse repositioning.

To determine airway flow (volume change/time) we first fitted the smoothed and drift-corrected volume signal with a cubic spline. Flow was obtained from the first-order time derivative (change in volume/change in time) of the cubic spline function.

We adapted the peak detection algorithm previously described by Noto *et. al* for locating the inspiratory peak and expiratory trough of each breath in humans (21). We adjusted the window sizes used by this algorithm from 300, 500, 700, 1000, and 5000 msec used in humans to 10, 20, 50, 100, and 200 msec for our mouse data to account for the faster breathing rate observed in mice compared to humans. The window shift sizes remained unchanged at 0%, 33%, and 66% of window size. Additionally, we included a smoothing step consisting of a Gaussian smoothing filter with a large sigma of 30 prior to the detection of consensus peaks and troughs. This reduced the detection of false-positive peaks and troughs caused by minor fluctuations in pressure.

To determine the presence of pauses in breathing between inspiratory and expiratory phases (end-inspiratory pause) we again adopted a previously described algorithm (21). Since we were primarily interested in pauses occurring between the inhale and exhale phases, and not those occurring midway through one of these phases, we limited the search for pauses to those that occurred within the first 25% of volume increase (expiration). For each breath, the airway flow signal between the breath’s peak and trough was used to identify the end of inhalation, the pause after inhalation, and the onset of exhalation.

Pauses were determined by creating a histogram of flow values from the peak to trough, where the range of flow values is equally split into 50 bins. We first identified the bin containing the zero-flow value. Then we examined the five bins on either side of the zeroth bin. Of these 11 bins, the bin with the highest count was identified as the center of the pause, or mode bin. A pause was detected if the mode bin count was at least two times larger than the mean count across all bins. Additional bins next to the mode bin were included as part of the pause if their count was at least 0.75 times that of the mode bin. In the case where no pause was detected, the end of exhalation and onset of inhalation were defined by the zero-crossing of the flow signal. When a pause was present, the end of inhalation and start of the post-inhale pause were defined as the point where the maximum flow value among pause bins was exceeded. Likewise, the end of the post-inhalation pause and start of exhalation were defined as the point where the minimum flow value among pause bins was exceeded.

Once the start and end of each phase of breathing were determined, the software identified the remaining important breathing landmarks and used those to calculate the variables that describe each breath (Table 1).

**Table 1.** Landmarks and Descriptive Variables Identified for Each Breath

| Landmarks |  |
| --- | --- |
| Landmark | Description |
| Inhale Onset | Start of inhale phase |
| Inhale Offset | End of inhale phase |
| Post-Inhale Pause Onset | Start of pause phase occurring after inhale phase |
| Post-Inhale Pause Offset | End of pause phase occurring after inhale phase |
| Exhale Onset | Start of exhale phase |
| Exhale Offset | End of exhale phase |
| Post-Exhale Pause Onset | Start of pause phase occurring after exhale phase |
| Post-Exhale Pause Offset | End of pause phase occurring after exhale phase |
| Inhale Flow Peak | Point of maximum flow during inhale and post-inhale pause phases |
| Exhale Flow Peak | Point of maximum flow during exhale and post-exhale pause phases |
| Inhale Volume Peak | Point of maximum volume during inhale and post-inhale pause phases |
| Exhale Volume Peak | Point of maximum volume during exhale and post-exhale pause phases |
| 50% Exhale Volume | Point where 50% of maximum volume during exhale phase is achieved |
| 64% Exhale Volume | Point where 64% of maximum volume during exhale phase is achieved |
| Descriptive Variables |  |
| Breath Measurement | Description |
| Inspiratory Volume | Volume change from inhalation onset to inhalation offset |
| Post-Inhalation Pause Volume | Volume change from post-inhalation pause onset to post-inhalation pause offset |
| Expiratory Volume | Volume change from exhalation onset to exhalation offset |
| Post-Exhalation Pause Volume | Volume change from post-exhalation pause onset to post-exhalation pause offset |
| Inhalation Peak Flow | Flow value at inhalation flow peak |
| Exhalation Peak Flow | Flow value at exhalation flow peak |
| Mid-Expiratory flow (EF50) | Flow value at 50% expiratory volume |
| Inspiratory Time | Time change from inhalation onset to inhalation offset |
| End-Inspiratory pause | Time change from post-inhalation pause onset to post-inhalation pause offset |
| Expiratory Time | Time change from exhalation onset to exhalation offset |
| Breath Duration | Time change from inhalation onset to post-exhalation pause offset |
| Estimated Breaths per Minute (BPM) | Estimated number of breaths per minute based on current breath's duration |
| Exhalation 64% Relaxation Time | Time change from exhalation onset to 64% exhale volume |
| Enhanced Pause (Penh) | $\frac{((Exhalation\ Duration)/(Exhalation\ 64\%\ Relaxation\ Time) - 1) \times ((Exhalation\ Peak\ Flow) / (Inhalation\ Peak\ Flow))}{1}$ |

Because mouse movement significantly alters the pressure signal, we had the software eliminate any measurements that were distorted using pre-defined, quality control algorithms. Any breaths where the original voltage signal exceeded minimum or maximum voltage saturation thresholds (0.150 and 4.800 Volts, respectively) were excluded. To determine if a breath coincided with mouse movement, the 50% inhale and exhale volume landmarks of each breath were compared to a movement threshold of 0.4 ml. For an ideal breath, both points should coincide with the global mean volume value. If the volume at either of these points was greater than 0.4 ml away from the global mean, the breath was excluded. Next, the inhale/exhale volumes, inhale/exhale peak flows, and inhale/exhale durations for each breath were compared to the mean values of these statistics for all other breaths within a 30-second window centered on that breath’s onset of inhalation. If any of these values were more than four standard deviations from the mean, the breath was excluded. A final control based on proximity to other excluded breaths was implemented, since measurement artifacts tended to span breathing segments that included multiple breaths. A breath was excluded based on proximity if it was located between two excluded breaths, and if it was less than four breaths away from each of these excluded breaths.

### Utilizing Software Programs for Analysis and Calibration

Differences in animal size alters free chamber volume and the relationship between change in volume and change in pressure. In turn, the chamber is calibrated each time an animal is placed in the chamber. The calibration curve was highly repeatable within and between mice with a within subject coefficient of variation of 5.38 ± 1.96% and between subject coefficient of variation of 5.55 ± 1.01% (Supplementary Table 1). Sensitivity limitations were assessed by identifying the residual difference between the known injected volume and calculated volume based on the regression curve. The average residual was 12.39 ± 0.134 µl resulting in a 95% confidence interval for sensitivity of 10.79 to 13.99 µl (Supplementary Table 1). In turn, 10.79 µl is the minimum change in breathing we can measure with confidence. Importantly, because this calculation was based on residuals from a known and calculated volume and our known volume (as measured using a 1 ml syringe) likely had error greater than 10 µl, this is likely an underestimate of our sensitivity using this system.

The software for calibration and analysis are a single program (https://github.com/bjrenquist/plethysmography; Figure 2B). We stopped the recording after calibration and restarted recording for the measurement meaning the calibration files saved separately from files we analyzed. For calibration files, the user opens the respiratory analysis program and opens the file with the calibration injections (clicking *File*◊*Open*◊*Select File*). When opened the user can see the voltage and pressure traces. To calibrate, the user goes to automatic calibration (*Analysis*→*Start Automatic Calibration*) and for calibration files selects *No* on the pop-up screen. Next, we visually examined the linear regression equation calculated by the program for a low number of points identified or a low R^2^ value. Before saving the calibration file, the time for the program to find breath landmarks (*Analysis*→*Find Breath Landmarks*) must be entered. The calibration file is then saved (*File*→*Save As*).

Once the corresponding calibration file was analyzed, the user opened the file to be analyzed. This time the user again selects automatic calibration (*Analysis*→*Start Automatic Calibration*) and selects *Yes* to use the calibration data from the calibration Microsoft Excel file previously saved. To complete the analysis, the user enters the time frame for the program to find breath landmarks and saves the output file in the appropriate folder. The output files can be opened in Excel for further analysis.

### Statistics

Mixed model analysis of variance was performed in SAS (SAS Inst., Cary, NC) for all analyses. To assess the dose response to methacholine we identified the maximal change from baseline in each mouse at each dose. Methacholine dose response was analyzed using a repeated measures ANOVA with dose as the main effect. Differences between means at each dose were corrected for multiple comparisons using a Tukey’s adjustment. The effect of HDM on tidal breathing was assessed using a repeated measures ANOVA with HDM exposure (pre- or post) as the main effect. The effect of HDM on the methacholine dose response was analyzed with a repeated measures ANOVA that included dose and HDM exposure as the main effects. Differences between means were investigated within a methacholine dose using Bonferroni correction for multiple comparisons. The effect of age (6 or 18 mo.) and sex (male or female) on tidal breathing was assessed using a two-way ANOVA. Differences between means were corrected for multiple comparisons using a Tukey’s adjustment.

## Results

### Chamber validation

To validate our system, we measured tidal breathing in lean adult male mice. We report inspiratory and expiratory volume, the time of inspiration and expiration, breaths per minute, enhanced pause (Penh), mid-expiratory flow (EF50), and end-inspiratory pause (also referred to as duration of braking) (17, 21). Inspiratory time, expiratory time, breaths per minute, Penh, and end-inspiratory pause measured by our system were all within published ranges (Supplementary Table 2).

We applied our head-out plethysmography system to measure breathing changes caused by acute bronchoconstriction induced by aerosolized methacholine (30 seconds of exposure) in sterile PBS (0, 25, 50 and 100 mg/ml at a nebulization rate of 0.191 ml/30 seconds). In line with previous reports, we found that methacholine dose-dependently decreased inspiratory/expiratory volume, EF50, and breaths per minute and increased inspiratory time, expiratory time, and Penh (Figure 3, Supplementary Table 3) (18, 26-30). Methacholine also increased end-inspiratory pause, a variable less commonly reported, but easily observable after methacholine administration (Figure 3O, P, Supplementary Figure 1 red in volume trace).

**Figure 3.**
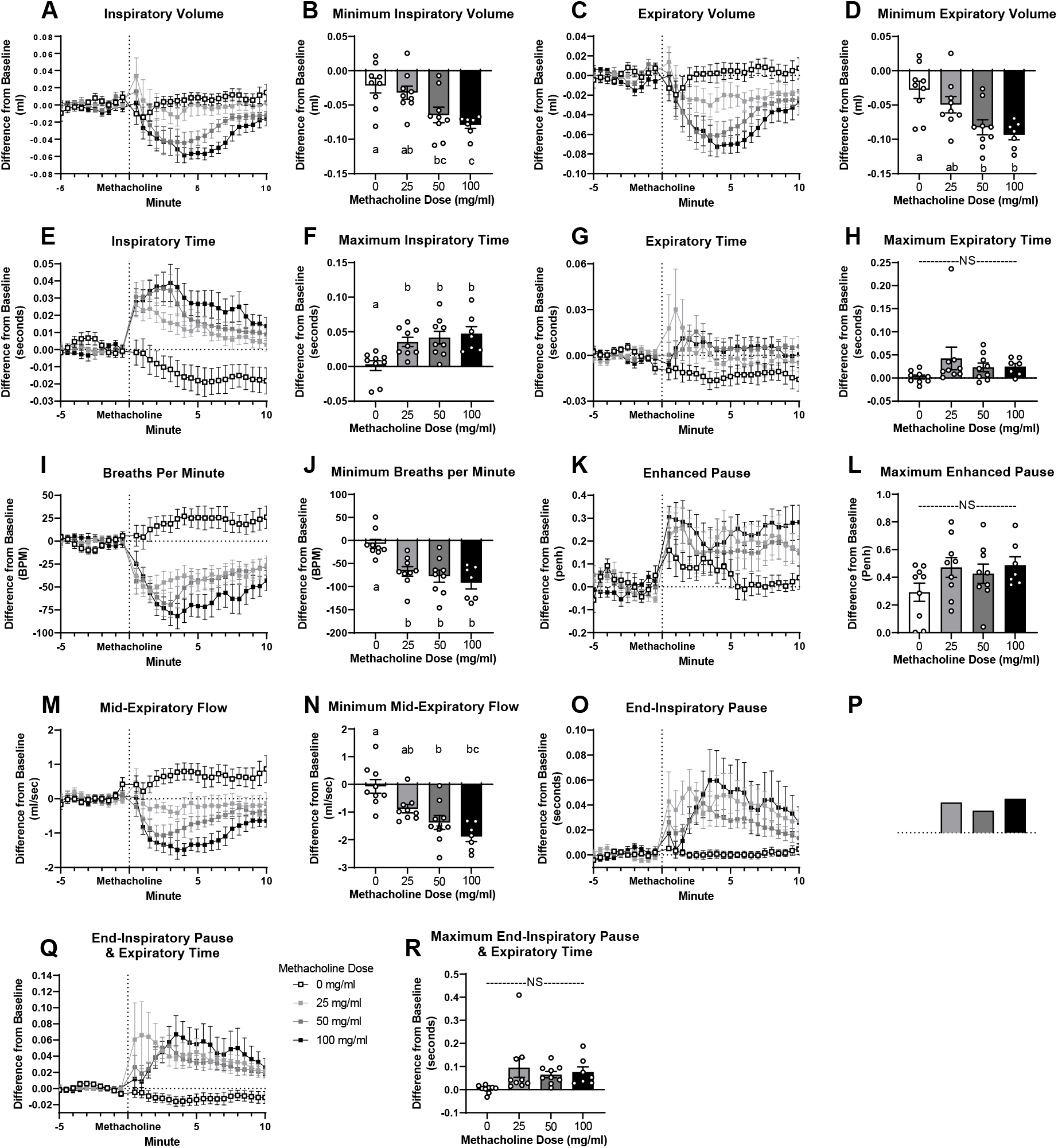
Response to aerosolized methacholine given at minute 0 (exposure 30 seconds; flow rate 0.191 ml/30 sec.). Figures A, C, E, G, I, K, M, O & Q report time course. Results presented as difference from a baseline measurement taken before methacholine exposure. Figures B, D, F, H, J, L, N, P & R report minimum/maximum 1 minute rolling average occurring within 10 minutes after methacholine administration. Data are presented as mean ± SEM. ^a,b,c^ bars that do not share a letter differ significantly (P < 0.05; n = 7-9).

We further validated the use of our system to measure alterations in lung function associated with HDM sensitization. HDM sensitization tended to decrease inspiratory volume (P = 0.067; Figure 4A) and Penh (P = 0.058; Figure 4F) and decreased expiratory volume (P = 0.016; Figure 4B), breaths per minute (P = 0.0093; Figure 4E), and EF50 (P = 0.015; Figure 4G). HDM sensitization also increased end-inspiratory pause (P = 0.0029; Figure 4H) and end-inspiratory pause and expiratory time (P = 0.014; Figure 4I). HDM sensitization increased the methacholine (100 mg/ml dose exposure for 30 seconds; ∼19.1 mg delivered total) induced increase in inspiratory time (P = 0.0156; Figure 5C) and end-inspiratory pause (P = 0.0508; Figure 5H) and tended to increase the methacholine extension of expiratory time (P = 0.0648; Figure 5B).

**Figure 4.**
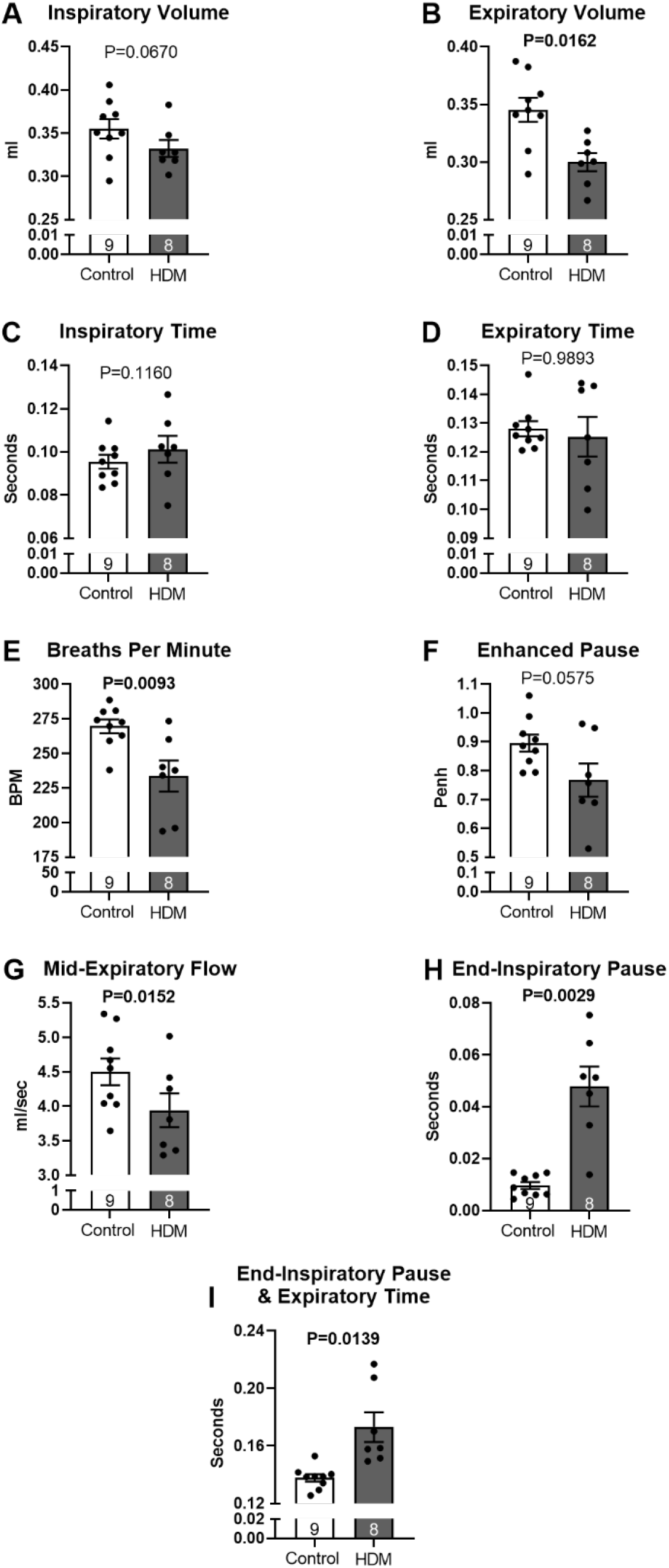
Tidal breathing in mice before (white) and after (gray) sensitization with house dust mite. Data are presented as mean ± SEM (n = 8-9).

**Figure 5.**
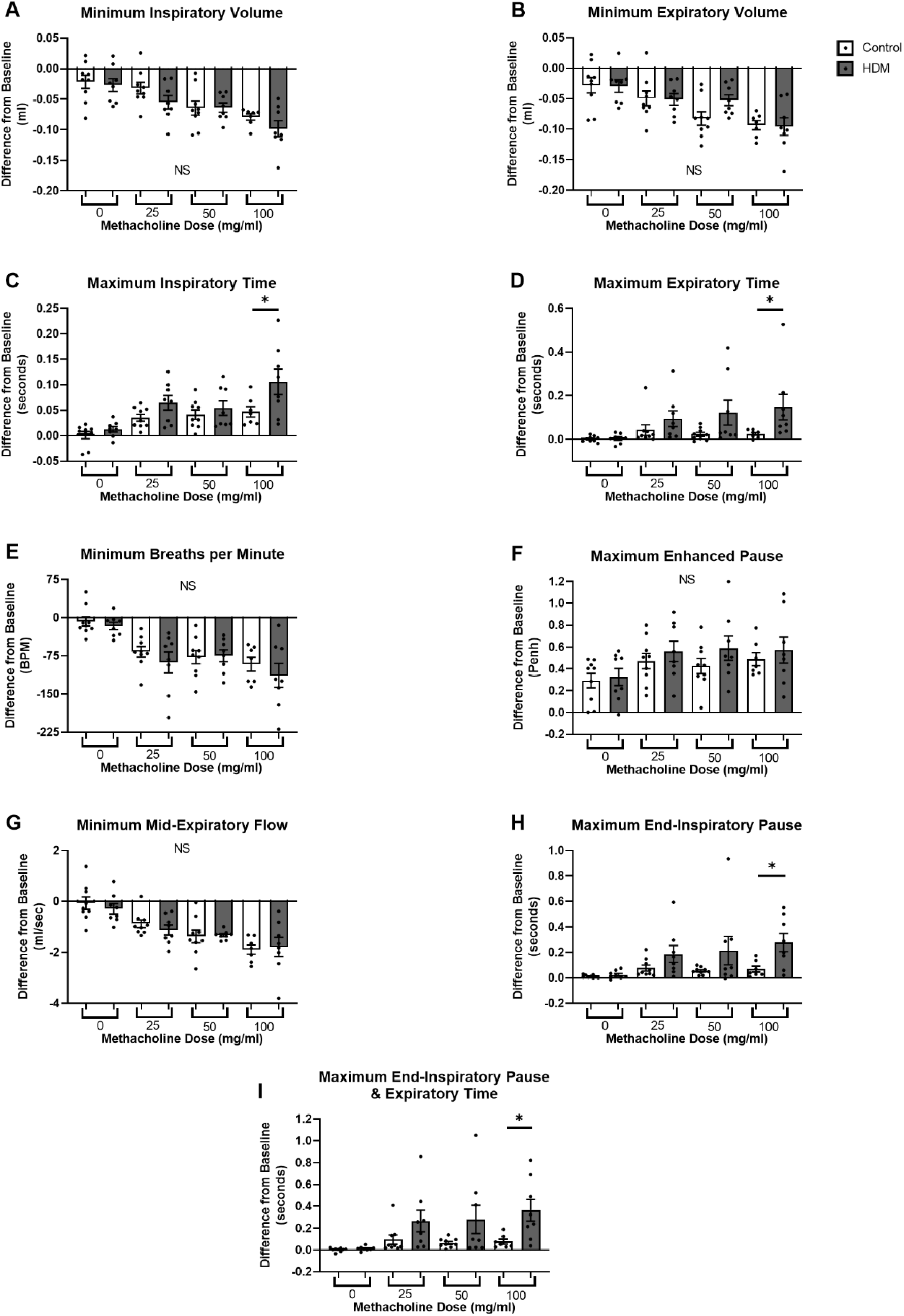
Minimal or maximal 1 minute rolling average over the 10 minutes after expose to methacholine (0, 25, 50, or 100 mg/ml exposure for 30 seconds nebulized at flow rate of 0.191 ml/30 sec) before and after sensitization with house dust mite. *Indicates significant differences (P < 0.05) before (white) and after (gray) house dust mite exposure. Data are presented as mean ± SEM (n = 8-9).

We further validated the use of our system to measure tidal breathing changes associated with sex and aging. Inspiratory (Male P = 0.0002; Female P = 0.0016; Figure 6A) and expiratory (Male P = 0.0026; Female P = 0.0102; Figure 6B) volume, inspiratory time (Male P = 0.0071; Female P = 0.0007; Figure 6C), and Penh (Male P < 0.0001; Female P = 0.0565; Figure 6F) increased with age in both male and female mice. Mid-expiratory flow increased with age in male mice (P = 0.0003; Figure 6G) and tended to increase with age in female mice (P = 0.11; Figure 6G). End-inspiratory pause (P = 0.016; Figure 6H) and end-inspiratory pause and expiratory time (P = 0.0176; Figure 6I) increased with age only in female mice. Breaths per minute decreased with age in both male and female mice (Male P = 0.0137; Female P = 0.0015; Figure 6E).

**Figure 6.**
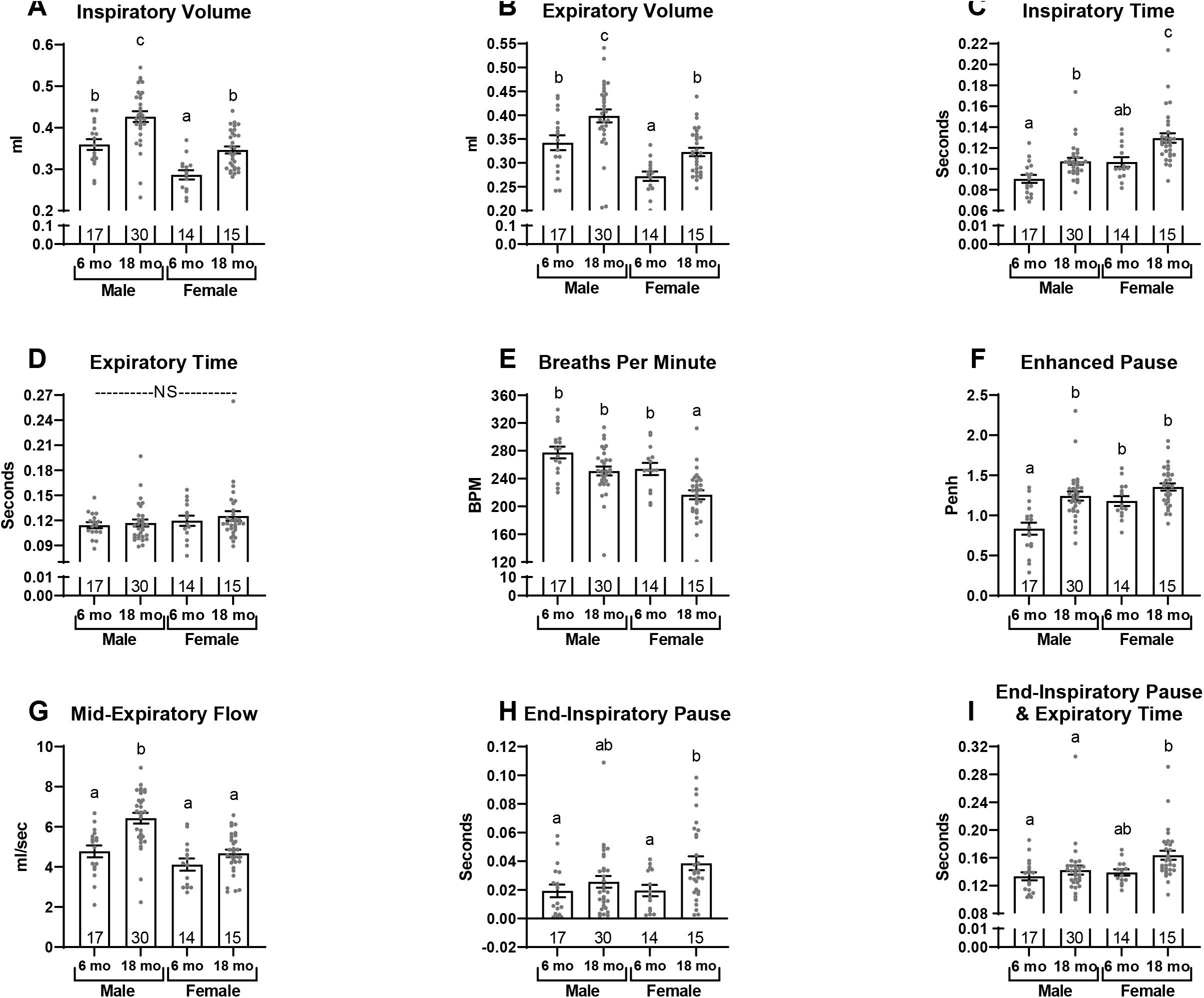
Tidal breathing in male and female mice aged 6 months (mo.) and 18 mo. Data are presented as mean ± SEM^. a,b,c^ bars that do not share a common letter differ significantly (P < 0.05; n =14-30).

## Discussion

By incorporating an inflatable balloon cuff that secures around the mouse’s neck, we created a leak free variable-pressure head-out plethysmography system that allows for accurate, repeatable measurements of volume and flow. Although inspiratory and expiratory volume are not direct measures of tidal volume, they are direct measures of the change in volume in the chamber resulting from expansion and contraction of the ribcage. Since air expands when warmed and humidified in the lungs, these measured values are larger than direct measures of tidal volume (Supplementary Table 2) (22).

HDM is commonly used to create a mouse model of asthma similar to the disease in humans (22, 23). Sensitization increases airway responsiveness, inflammation, and induces airway remodeling (24-26). We measured alterations in tidal breathing, including decreases in volume and EF50, consistent with reports of tidal breathing in humans with asthma and HDM sensitized mice (27-30). Individuals with asthma have increased respiratory rate, inspiratory flow rate, and decreased minute ventilation, tidal volume, and inspiratory time (27-29). Airway hyper-responsiveness to methacholine is characteristic of asthma (31). HDM sensitization in mice increases the responsiveness to methacholine, with greater decreases in EF50, increases in Penh, and increases in airway resistance and elastance (18, 32-34). Herein, we measure HDM sensitization increases methacholine induced changes in inspiratory and expiratory time and the end-inspiratory pause indicative of the expected increased airway-hyperresponsiveness in a HDM sensitized mouse.

The variable most reported from whole-body plethysmography measurements is Penh, a dimensionless variable designed to describe the waveform of each breath (3, 33). Penh is mouse strain specific and may not correlate with more sensitive measures of lung function, such as those measured using forced oscillation technique (1, 27, 34). Though we did measure an increase in Penh following administration of a bronchoconstrictor (Figure 3K, L), we recognize that Penh may not correlate with the gold-standard invasive measurements of resistance, tissue damping, and airway elastance (35).

This head-out plethysmography system does not measure resistance, compliance, conductance, or elastance which are direct, sensitive measures of lung mechanics that serve as the gold standard for rodent lung function. With minor modifications this system could be adjusted to become a double-chamber plethysmography system allowing for the measurement of specific airway resistance and specific airway conductance. Our design incorporating a leak-proof body chamber would eliminate the potential for air leak between the nasal and thoracic chamber.

Previous studies using forced oscillation technique report that methacholine increases airway resistance, airway elastance, tissue damping, and tissue elastance and decreases conductance and compliance (23-25). We report methacholine induced changes in breaths per minute, EF50, expiratory time, and volume, all variables which are associated with increases in airway resistance, and changes in conductance and dynamic compliance (14, 21, 27-29). We have further shown that inspiratory/expiratory volume, breaths per minute, EF50, and end-inspiratory pause are sensitive to aging, corresponding to age-related alterations in resistance and compliance (31, 32).

With this system, animals can recover from previous exposure to a bronchoconstrictor, or other treatment, before receiving the next dose. We have shown that many variables assessed with head-out plethysmography (expiratory/inspiratory volume, inspiratory time, breaths per minute, and end-inspiratory pause) do not return to baseline within 10 minutes after exposure to a low dose of methacholine (25 mg/ml exposure for 30 seconds; total of ∼4.775 mg methacholine) (Figure 3A, C, E, I, O). Without recovery bronchoconstriction from a previous dose may limit subsequent delivery of methacholine attenuating the effect of incremental increases in methacholine dose (35, 36). Delivery of methacholine is not limited in studies applying forced oscillation technique, as deep inflation, inflating the lungs to total lung capacity, is performed after each methacholine dose (35). However, the reported response to methacholine using whole-body plethysmography uses an incremental protocol without allowing for sufficient time for mice to recover (37-39). Finally, utilizing our head-out plethysmography system, we can observe the time course of response to an acute intervention rather than being limited to reporting a maximal response (Figure 3) (33, 39-42). Many investigations only present a brief period of normal breathing (e.g. 10 breaths or 45 seconds) and methacholine response as the maximum or minimum response (10, 18, 35, 39, 41, 43). We designed our software to automatically calculate rolling averages over a long timeframe to ensure a more accurate time-dependent model.

We report greater amounts of bronchoconstriction (both percent change and minimum/maximum percent change) at lower methacholine doses than previously reported in studies using whole-body plethysmography (gray Supplementary Table 3). Importantly, to avoid potential confounding effects of prior bronchoconstriction, we only compared our results to the first dose given in previous studies (white Supplementary Table 3). In addition, although we use C56BL/6 mice, which have a diminished response to methacholine, we report greater changes in volume, inspiratory time, breaths per minute, Penh, and EF50 at lower doses of methacholine (Supplementary Table 3) (35, 44). Our enhanced response may be due to improved drug delivery, better software capturing data for each breath, and/or enhanced system sensitivity. We aerosolized methacholine directly onto the face of the mouse instead of into a whole-body plethysmography chamber. In addition, we utilized a nebulizer that produces small particle size (4.0-6.0 µm) which are more easily delivered further into the lung. Finally, our airtight chamber allows for accurate determination of changes in volume and flow.

Forced oscillation technique, the gold standard, cannot be used to assess the respiratory drive of breathing, which can only be measured in an awake, conscious mouse. Control of respiration is altered in asthma, chronic obstructive pulmonary disorder, morbid obesity, or neuromuscular diseases (45-48). The respiratory drive of breathing is also altered in mouse models of disease, such as asthma or obesity, and under hypoxic and hypercapnic conditions which can be assessed by measuring the volume and the timing of each breath (33, 49, 50).

We developed and validated a variable-pressure head-out plethysmography system that can accurately assess breath volume and timing across a large timeframe, with multiple replicates in the same mouse. We created corresponding software to collect and analyze large datasets. This provides an attractive alternative to forced oscillation technique, whole-body plethysmography, and commercially available head-out plethysmography systems to study pulmonary function and disease in mice.

## Acknowledgements

We want to acknowledge the researchers that helped with analysis of the data: Skylar Knight, Kendra Miller, and Emily Ngu. Finally, we would also like to thank Dr. Janko Nikolich-Žugich and Jennifer Uhrlaub for the use of mice from their colony.

---

**Supplementary Figure 1.**
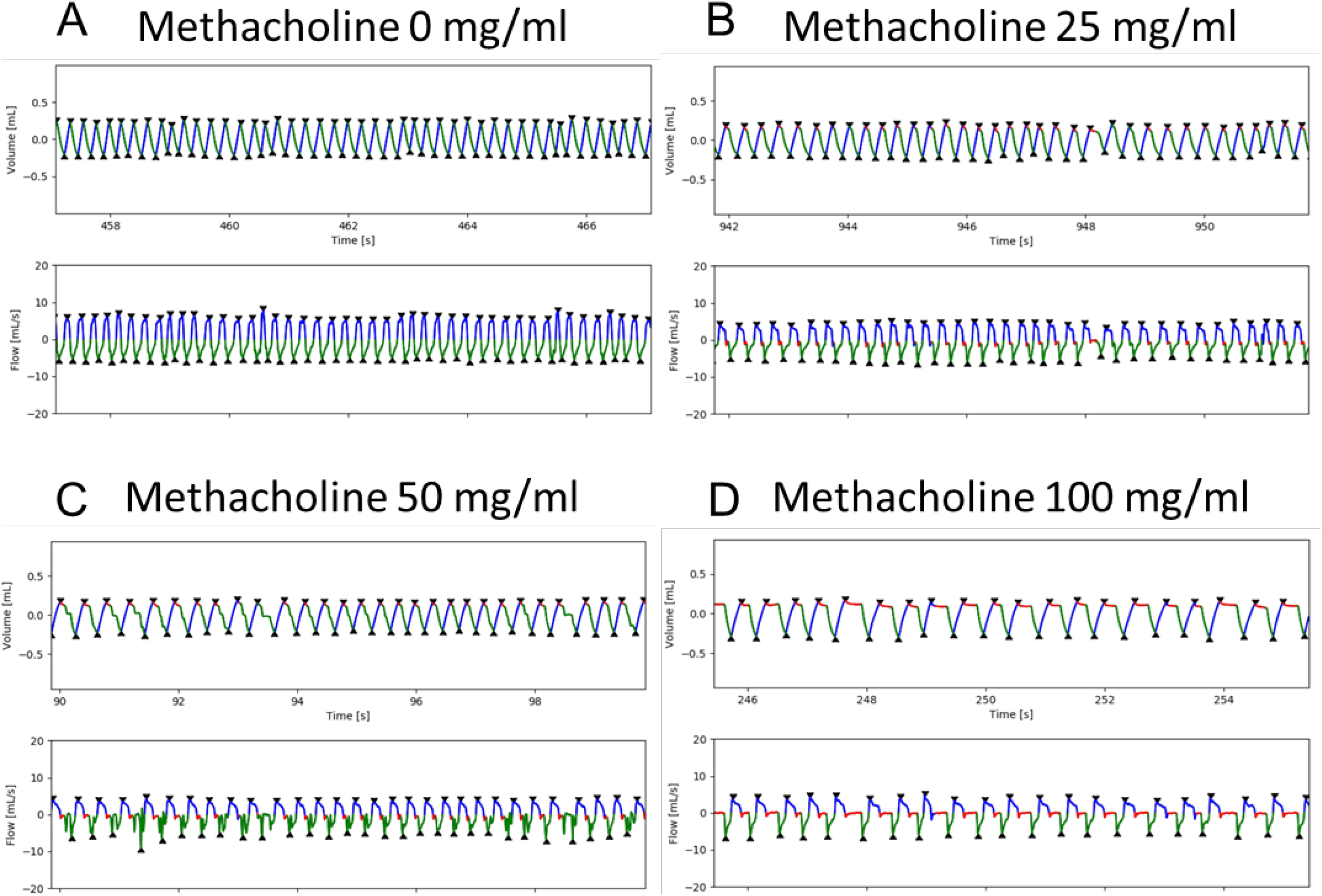
Representative sample of breathing trace after exposure to bronchoconstrictor A) 0 B) 25 C) 50 D) 100 mg/ml of methacholine for 30 seconds at flow rate of 0.191 ml/30 sec. Blue = inspiration, green = expiration, red = end-inspiratory pause.

**Supplementary Table 1.** Repeated calibrations in the same mouse to calculate variability within and across mice and residual difference between calculated and actual change in volume during calibrations to identify the sensitivity of measurement.

| Mouse | Mouse Weight (g) | Session Number | Regression Equation | R<sup>2</sup> Value | Average Residual Value (μl) | SEM |
| --- | --- | --- | --- | --- | --- | --- |
| 1 | 23.4 | 1 | y=1318.43x | 0.99 | 25.27 | 0.0014 |
| 1 |  | 2 | y=1207.74x | 0.93 | 14.59 | 0.0009 |
| 1 |  | 3 | y=1272.37x | 0.95 | 13.55 | 0.0008 |
| 1 |  | 4 | y=1338.53x | 0.99 | 14.42 | 0.0009 |
| 2 | 21.5 | 1 | y=1179.82x | 0.99 | 12.98 | 0.0009 |
| 2 |  | 2 | y=1319.87x | 0.99 | 21.55 | 0.0018 |
| 2 |  | 3 | y=1314.65x | 0.99 | 22.36 | 0.0024 |
| 2 |  | 4 | y=1411.60x | 0.99 | 9.29 | 0.0009 |
| 3 | 22.6 | 1 | y=1200.79x | 0.98 | 9.29 | 0.0007 |
| 3 |  | 2 | y=1361.35x | 1.00 | 11.05 | 0.0006 |
| 3 |  | 3 | y=1325.56x | 0.99 | 8.24 | 0.0005 |
| 3 |  | 4 | y=1206.01x | 0.99 | 12.08 | 0.0008 |
| 4 | 22.3 | 1 | y=1378.43x | 1.00 | 10.82 | 0.0012 |
| 4 |  | 2 | y=1199.40x | 0.95 | 4.73 | 0.0004 |
| 4 |  | 3 | y=1180.05x | 0.95 | 10.78 | 0.0010 |
| 4 |  | 4 | y=1262.16x | 0.99 | 10.41 | 0.0007 |
| 5 | 20.2 | 1 | y=1131.72x | 0.99 | 5.09 | 0.0003 |
| 5 |  | 2 | y=1245.60x | 0.99 | 22.01 | 0.0015 |
| 5 |  | 3 | y=1153.74x | 0.98 | 12.04 | 0.0008 |
| 5 |  | 4 | y=1248.37x | 0.97 | 8.04 | 0.0006 |
| 6 | 19.8 | 1 | y=1223.48x | 0.98 | 12.32 | 0.0010 |
| 6 |  | 2 | y=1314.37x | 0.98 | 10.98 | 0.0006 |
| 6 |  | 3 | y=1356.95x | 0.98 | 10.37 | 0.0008 |
| 6 |  | 4 | y=1230.22x | 0.98 | 15.38 | 0.0010 |
| 7 | 21.5 | 1 | y=1140.25x | 0.99 | 14.24 | 0.0007 |
| 7 |  | 2 | y=1356.88x | 0.98 | 14.09 | 0.0006 |
| 7 |  | 3 | y=1223.15x | 1.00 | 14.65 | 0.0006 |
| 7 |  | 4 | y=1303.98x | 0.97 | 14.55 | 0.0009 |
| 8 | 17.7 | 1 | y=1162.83x | 0.97 | 9.79 | 0.0009 |
| 8 |  | 2 | y=1266.71x | 1.00 | 13.05 | 0.0012 |
| 8 |  | 3 | y=1204.77x | 0.99 | 6.55 | 0.0004 |
| 8 |  | 4 | y=1243.15x | 1.00 | 18.41 | 0.0010 |
| 9 | 18.2 | 1 | y=1278.32x | 0.96 | 13.37 | 0.0010 |
| 9 |  | 2 | y=1242.87x | 0.98 | 6.81 | 0.0005 |
| 9 |  | 3 | y=1252.19x | 0.99 | 7.66 | 0.0004 |
| 9 |  | 4 | y=1283.92x | 0.98 | 5.48 | 0.0003 |

**Supplementary Table 2.**
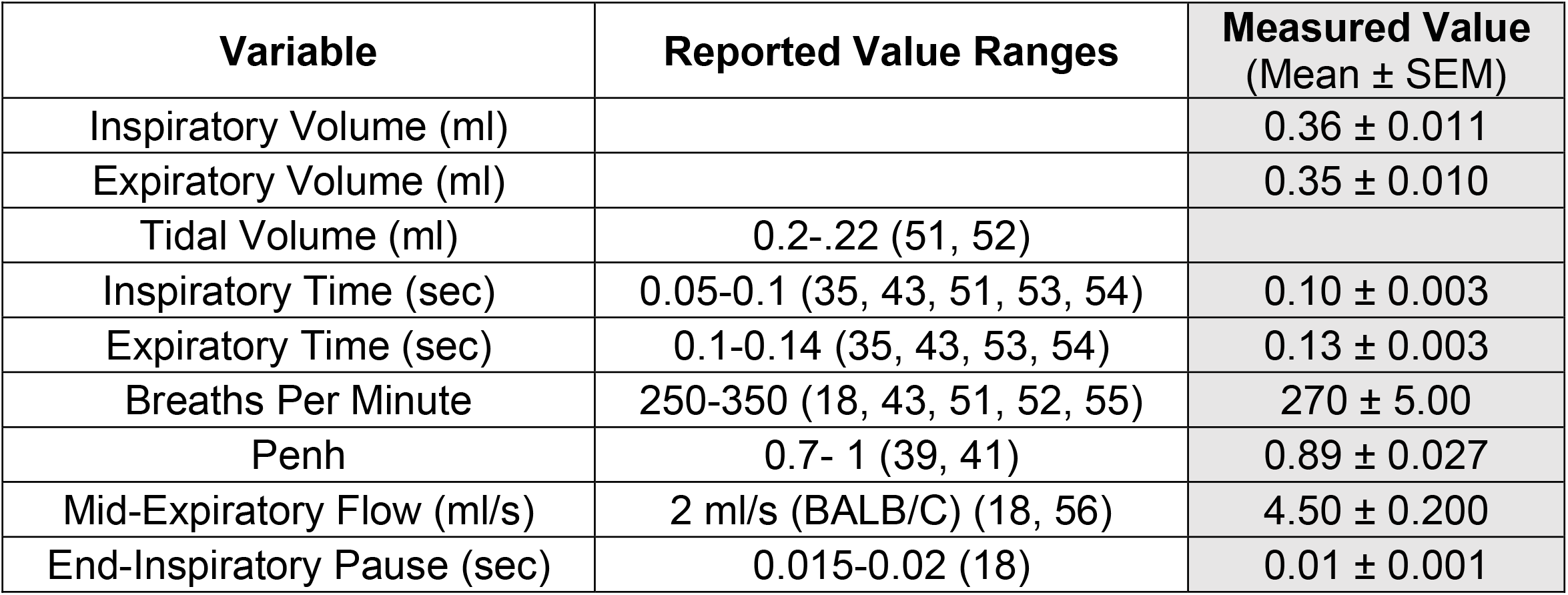
Tidal Breathing Validation

**Supplementary Table 3.** Comparison of Methacholine Response Between Studies (Whole-Body Plethysmography)

| Post Methacholine 5 Minute Average Percent Change from Baseline |  |  |  |  |  |  |  |
| --- | --- | --- | --- | --- | --- | --- | --- |
| Variable | Percent Change (%) | Methacholine Dose (mg) | Mouse & Sex | Reference | Percent Change (%) | Methacholine Dose (mg) | Mouse & Sex |
| Tidal Volume | 6.04 | 19.1 | Unspecified | (10) | -12.56 | 19.1 | Male C57BL/6 |
|  | -5.17 | 4700 | Female C57BL/6 | (35) |  |  |  |
| Inspiratory Time | 1.85 | 19.1 | Unspecified | (10) | 37.52 |  |  |
| Expiratory Time | 12.79 | 19.1 | Unspecified | (10) | 4.54 |  |  |
| Breaths Per Minute | -7.07 | 19.1 | Unspecified | (10) | -22.17 |  |  |
|  | -11.59 | 4700 | Female C57BL/6 | (35) |  |  |  |
| Penh | -3.21 | 19.1 | Unspecified | (10) | 28.79 |  |  |
| Post Methacholine 2 Minute Average Percent Change from Baseline |  |  |  |  |  |  |  |
| Expiratory Time | 50 | 5700 | Female C57BL/6 | (27) | 4.54 | 19.1 | Male C57BL/6 |
| Penh | 0 | 5700 | Female C57BL/6 | (27) | 28.79 |  |  |
| Post Methacholine Minimum/Maximum Value within 3 Minutes Percent Change from Baseline |  |  |  |  |  |  |  |
| Penh | 8.33 | 6 | Male C57BL/6 | (37) | 31.78 | 19.1 | Male C57BL/6 |
| EF50 | -1.67 | 0.239 | Unspecified Balb/c | (18) | -15.63 |  |  |
| White = Values Reported in Other Studies; Gray = Author's Reported Values |  |  |  |  |  |  |  |

